# A combination of Metformin and Epigallocatechin Gallate Potentiates Glioma Chemotherapy *in vivo*

**DOI:** 10.1101/2022.11.16.516766

**Authors:** Shreyas S Kuduvalli, S Daisy Precilla, Anandraj Vaithy, Mugilarasi Purushothaman, Arumugam Ramachandran Muralidharan, B Agiesh Kumar, Markus Mezger, Justin S Antony, Madhu Subramani, Biswajit Dubashi, Indrani Biswas, K P Guruprasad, T.S Anitha

## Abstract

Glioma is the most devastating high-grade tumor of the central nervous system, with dismal prognosis. Existing treatment modality does not provide substantial benefit to patients and demands novel strategies. One of the first-line treatments for glioma, temozolomide, provides marginal benefit to glioma patients. Repurposing of existing non-cancer drugs to treat oncology patients is gaining momentum in recent years. In this study, we investigated the therapeutic benefits of combining three repurposed drugs, namely, metformin (anti-diabetic) and epigallocatechin gallate (green tea-derived antioxidant) together with temozolomide in a glioma-induced xenograft rat model. Our triple-drug combination therapy significantly inhibited tumor growth *in vivo* and increased the survival rate (50%) of rats when compared with individual or dual treatments. Molecular and cellular analyses revealed that our triple-drug cocktail treatment inhibited glioma tumor growth in rat model through ROS-mediated inactivation of PI3K/AKT/mTOR pathway, arrest of the cell cycle at G1 phase and induction of molecular mechanisms of caspases-dependent apoptosis. In addition, the docking analysis and quantum mechanics studies performed here hypothesize that the effect of triple-drug combination could have been attributed by their difference in molecular interactions, that maybe due to varying electrostatic potential. Thus, repurposing metformin and epigallocatechin gallate and concurrent administration with temozolomide would serve as a prospective therapy in glioma patients.

## 1. Introduction

Glioma accounting for 60% of all the adult brain tumors, is one among the deadliest central nervous system (CNS) malignancies worldwide (Jemal et al., 2010). Despite advanced surgical procedures, followed by radio-sensitization and treatment with temozolomide (T), a first-line chemotherapy drug, the median survival rate remains low in glioma patients with a 5 -year survival rate of less than 5% (Tamimi and Juweid, 2017). Disappointingly, the tumor cells have developed chemo-resistance against T, which calls out for effective therapeutic interventions and combination regimens to alleviate glioma pathogenesis (Lee, 2016). In the pipeline of targeted therapies against glioma, drugs that target signalling pathways responsible for tumor cell growth and progression have been identified to play vital role in alleviating glioma aggressiveness (Chowdhury et al., 2018). These drugs, when used either alone or in combination with the existing therapy, may progress the treatment outcome for individuals identified with this devastating tumor.

Reactive oxygen species (ROS), primarily produced in the mitochondria, contribute to the pathogenesis of various solid tumors including glioma (Rinaldi et al., 2016). Previous literature have proved that glioma tumor cells has the potency to survive under highly -stressed micro-environment (De Vleeschouwer and Bergers, 2017). This makes them more vulnerable to the damage caused by anti-tumor drugs, that are known to induce or enhance oxidative stress such as camptothecin, epigallocatechin gallate (E), kaempferol, chloroquine, etc. (Mileo and Miccadei, 2016; Sharma et al., 2007; Vessoni et al., 2016; Xia et al., 2005). Also, ROS regulate several signalling pathways such as phosphoinositide 3-kinase /protein kinase B/mammalian target of rapamycin (PI3K/AKT/mTOR) pathway and mitogen-activated protein kinase (MAPK) pathway (Koundouros and Poulogiannis, 2018). Among those, PI3K/AKT/mTOR pathway is a critical pro-survival signalling pathway, that modulates cell growth, proliferation, apoptosis and survival in tumor cells (Gravina et al., 2019). Aberrant activation of PI3K/AKT/mTOR pathway is observed in 80% of glioma cases, thereby emphasizing its significance as a therapeutic target (Mao et al., 2012). Nevertheless, the activation of PI3K/AKT/mTOR pathway in glioma leads to the development of drug resistance, thereby inhibiting the therapeutic effect of T (Li et al., 2016; Xia et al., 2020). It has also been reported that in most malignancies including glioma, the apoptotic pathway is usually inactivated by the activation of various pathways such as PI3K/AKT/mTOR, MAPK, Nuclear factor erythroid 2-related factor 2 (Nrf2) to maintain tumorigenesis (Redza-Dutordoir and Averill-Bates, 2016). These pathways mainly suppress the expression of key activators of the apoptotic pathway such as the BCL2 associated X and agonist of cell death (BAX/BAD) complex and p53 activation. Meanwhile, a previous study demonstrated that augmentation of ROS can inactivate PI3K/AKT/mTOR pathway and may promote apoptosis (Usatyuk et al., 2014). Hence, unravelling the multi -faceted interactions between ROS metabolism and PI3K/AKT/mTOR, paving a way out for apoptosis will undoubtedly provide novel insights on the identification of exploitable vulnerabilities for treatment of hyperactive PI3K/AKT/mTOR tumors like glioma.

Underpinning the above ideas, recent studies have shown that metformin (M), a major anti-diabetic drug, has exhibited synergistic effects with T treatment and has the potential to enhance its chemotherapeutic efficacy in glioma, thereby opening a new avenue to overcome T resistance in glioma therapy (Samuel et al., 2019). Recently, Valtorta et al., has documented that the synergism of M with T reduced oxidative stress in U251 and T98G cell lines (Valtorta et al., 2017). Further, individually M has been proven to bring about 5’ adenosine monophosphate-activated protein kinase (AMPK) mediated inhibition of fork-head box O3 (FOXO3) and AKT in various cancers (Chou et al., 2014; Mazurek et al., 2020; Nozhat et al., 2018; Sato et al., 2012). E, one of the major flavonoids in green tea has been bestowed with immense anti-oxidant activity (25-100 times higher) than vitamin-C and vitamin-E, and is considered to be one of the dietary therapeutic potential compounds (Thawonsuwan et al., 2010). Studies have also suggested E as a potential chemotherapeutic agent, as it has shown effective inhibitory effects on generation of ROS, cell proliferation (Almatroodi et al., 2020; Du et al., 2012) and on key proteins involved in cell cycle regulation in glioma (Cheng et al., 2020). Interestingly, a study by Zhang et al., showed that E individually enhanced the anti-glioma efficacy of T in U87 human glioma cell line (Zhang et al., 2015). However, so far only a single study, has evaluated the combined efficacy of M and E to suppress Nrf2 pathway, in non-small cell lung cancer, but none has been reported in glioma (Yu et al., 2017).

To add-on, the major restraint in the use of T to treat glioma is that, it induces haematotoxicity, leading to a clinically significant toxicity and development of chemo-resistance (Fernandes et al., 2017). To overcome these issues, the combined treatment of a synthetic compound and an antioxidant by means of repurposing paradigm, could be a better approach to prevent glioma growth and proliferation. Repurposing of anti-diabetic drug, M and an antioxidant, E in glioma treatment together with T has several advantages including early clinical adoption, cost-effectiveness, proved drug safety and availability of human pharmacological data. Hence, in the present study, we elucidated the anti-glioma efficacy of the triple-drug combination (TME) in a glioma-induced xenograft model. In addition, the relationship between ROS and PI3K/AKT/mTOR pathway leading to apoptosis following the treatment with this triple-drug combination was also investigated in glioma-induced rats.

## 2. Methods

### 2.1 Chemicals

Minimum Essential Medium Eagle (EMEM) and penicillin-streptomycin were procured from HiMedia (Mumbai, India) while fetal bovine serum (FBS) was obtained from Gibco Thermo Fisher (Massachusetts, USA). T (Cat No: 85622-93-1), M (Cat No: 1115-70-4) and E (Cat No: 989-51-5) were obtained from Sigma Chemical Co (St. Louis, Missouri, USA). All other chemicals and reagents were of analytical or molecular grade and were obtained from HiMedia and Sigma Chemical Co.

### 2.2 Cell Culture

C6 rat glioma cell line (Passage no. 58) was procured from National Centre for Cell Sciences (NCCS), Pune, India and were employed for establishing the orthotopic xenograft glioma tumor. C6 cells were cultured in T75 flasks containing EMEM medium supplemented with 10% FBS and 1% penicillin-streptomycin. The cultures were maintained at 37^0^C in a humidified atmosphere containing 5% CO_2_.

### 2.3 Animals

Healthy male Wistar rats (180–240g / 6 to 8 weeks old) were purchased from Manipal Academy of Higher Education (MAHE), Manipal, India. Animals were maintained in a ventilated and temperature-controlled atmosphere at 23–25^0^C with a 12 h light/dark cycle and had access standard food pellets and water. All animals used in the present study received utmost humane care, and all experimental procedures were performed in accordance with institutional guidelines and regulations of Sri Balaji Vidyapeeth (Deemed to-be University), Puducherry, India. All the animal experimental protocols were reviewed, approved and performed in accordance with the guidelines and regulations set by the Institutional Animal Ethics Committee (IAEC), Kasturba Medical College, Manipal Academy of Higher Education (MAHE), Manipal (IAEC/KMC/69/2019) and Mahatma Gandhi Medical College and Research Institute, Sri Balaji Vidyapeeth (Deemed to-be University), Puducherry (07/IAEC/MG/08/2019-II). All the experiments were conducted in accordance to ARRIVE guidelines.

### 2.4 Orthotopic xenograft glioma model

Wistar rats were anesthetized with a ketamine/xylazine cocktail solution (87 mg/kg body weight and 13 mg/kg body weight) and were placed in the stereotaxic head frame. A 1-cm midline scalp incision was made, and 1×10^6^ C6 rat glioma cells in a volume of 3 μl phosphate buffer saline (PBS) was injected at a depth of 6.0 mm in the right striatum (coordinates with regard to Bregma: 0.5 mm posterior and 3.0 mm lateral) through a burr hole in the skull using a 10-μl Hamilton syringe to deliver tumor cells to a 3.5-mm intra-parenchymal depth. The burr hole in the skull was sealed with bone wax and the incision was closed using dermabond. The rats were monitored daily for signs of distress and death.

### 2.5 Study Design

This study was conducted on nine series of experimental groups (n=48), namely Group I: Normal rats without tumor induction (Control C) (n=3); Group II: Tumor-control (TC) (n=3); Group III: T-treated glioma-induced rats (T) (n=6); Group IV: M-treated glioma-induced rats (M) (n=6); Group V: E-treated glioma-induced rats (E) (n=6); Group VI: TM-treated glioma-induced rats (TM) (n=6); Group VII: TE-treated glioma-induced rats (TE) (n=6); Group VIII: ME-treated glioma-induced rats (ME) (n=6) and Group IX: TME-treated glioma-induced rats (TME) (n=6) as shown in **Figure 1A**. The dosage of the drugs, either alone or in combination, to be administered to the rats were selected based on our *in vitro* data and previous studies in glioma (Kuduvalli et al., 2021; Fernandes et al., 2017; Yu et al., 2017; Piwowarczyk et al., 2020).

**Figure 1.**
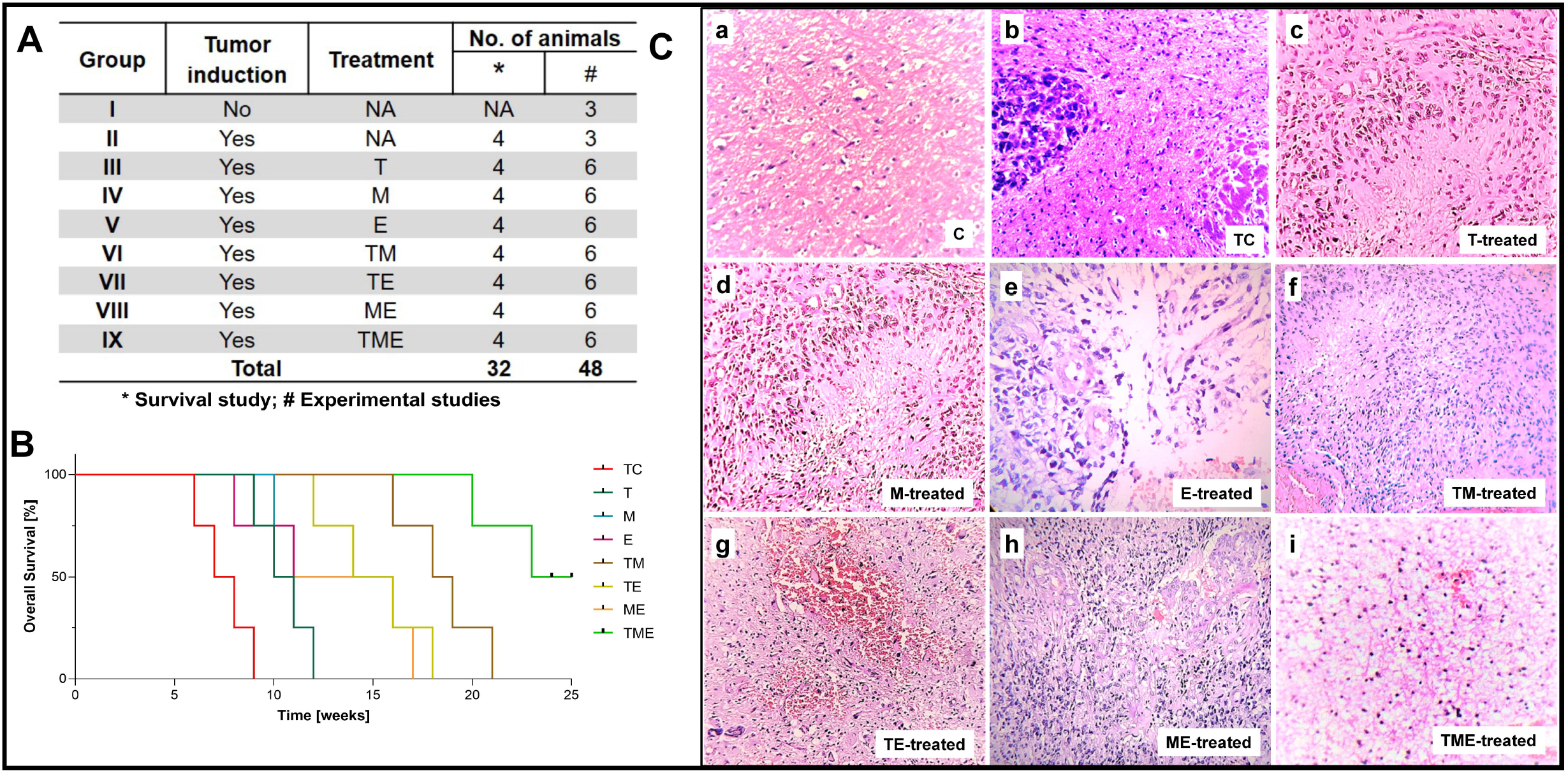
(**A**) Table showing the treatment strategies followed in the present study (* Survival study; # Experimental studies). (**B**) Kaplan-Meier survival curves were plotted and statistical analysis (Wilcoxon-Gehan test) was performed. (**C**) H & E staining of tumor sections from various treatment groups. Images were captured at 20x magnification, wherein the images C-a and C-b correspond to C and TC, while C-c to C-i correspond to treatment with T, M, E, TM, TE, ME and TME.

Treatment with the drugs was commenced 20 days after orthotopic implantation of the human glioma cells. As our *in vitro* data did not reveal any significant changes in the vehicle-treated cells when compared with glioma cells, the vehicle-treated rats were not further considered for the *in vivo* studies. After 7 days of treatment with the drugs, animals were anesthetized and 3-4 ml of blood was collected by cardiac puncture, using a 5 ml syringe. These animals were then sacrificed by euthanizing with isosulfan followed by cervical dislocation and the brain tissue was isolated, stored accordingly and were subjected to further studies.

### 2.6 Drug preparations

T, M and E, individually and in combination were prepared on each day of injection in sterile water (vehicle) at varying concentrations as listed in **Table 1,** respectively. The drugs were stored at 4^0^C before administration and were injected within 1 hour of formulation. All the drugs were administered intra-peritoneally in a volume of 0.1 ml.

**Table 1:** The concentration of each drug used in the treatment of animals both individually and in combination.

| Groups | Drugs | Dosage (mg/kg body weight) |
| --- | --- | --- |
| Group II | Tumor control | NA |
| Group III | T | 10 |
| Group IV | M | 15 |
| Group V | E | 25 |
| Group VI | TM | 10+15 |
| Group VII | TE | 10+25 |
| Group VIII | ME | 15+25 |
| Group IX | TME | 10+15+25 |

### 2.7 Survival statistics and toxicity studies

Survival study was carried out on a new set of thirty-two (N=32) rats that were implanted with C6 human glioma cells and treated with the respective drugs as mentioned previously in **Table 1**. After the treatment period, the animals were continuously monitored for their survival rate for a period of 25 weeks. In the survival analysis, the death of a rat in each group of treatment was taken as a break point and a survival graph was plotted using GraphPad Prism 8 (San Diego, USA). Any animal surviving a period of 25 weeks was euthanized under strict ethical guidelines, to avoid further suffering of the animal. Postmortem, drug-induced toxicity was determined by histopathological analysis of major organs namely liver, lungs, spleen, kidney and pancreas.

### 2.8 Haematoxylin and Eosin staining

After treatment for 7 days, the rats were euthanized and the brain tissues of the experimental groups were collected and fixed in 4% PBS-buffered paraformaldehyde followed by embedding in paraffin. In order to perform the hematoxylin and eosin (H & E) staining, paraffin blocks were sectioned by 5 μm thickness. Slides were then stained with H & E and were later observed under 20x magnification in a compound microscope (Axiovert, Carl Zeiss, USA) to take digital photographs.

### 2.9 Immunohistochemistry

For immunohistochemistry (IHC) analysis, 4 μm thick tissue sections were deparaffinized in xylene and hydrated by immersing in a series of graded ethanol concentration (100%, 95%, 75% and 50%). Endogenous peroxidase activity was blocked by incubating sections in 3% H_2_O_2_ solution prepared in methanol at room temperature for 10 minutes and were washed with PBS twice for 5 minutes each. Antigen retrieval was performed by immersing the slides in 300 ml of retrieval buffer (10 mM citrate buffer, pH 6.0) for one hour at 100^0^C and were rinsed twice in PBS. Sections were then incubated with approximately 100 μl diluted primary antibodies of Ki67 (1:100 dilution, Elabscience, Wuhan, China) and Vascular Endothelial Growth Factor (VEGF) (1:100 dilution, Elabscience, Wuhan, China) for 30 minutes at room temperature. The slides were then washed twice with PBS and were then incubated with approximately 100 μl of diluted secondary antibody (Cat No. E-IR-R213, Elabscience, Wuhan, China) at room temperature for 30 minutes. Slides were then washed in PBS for 5 minutes, followed by incubation with approximately 100 μl of diluted Horse Radish Peroxidase (HRP) conjugate for 30 minutes. The slides were then incubated in 100 μl diluted 3,3’-Diaminobenzidine (DAB) chromogen substrate for the development of color and the sections were counterstained by hematoxylin for 1-2 minutes. Later, the slides were washed under running tap water to remove excess stain and dried at room temperature. The sections were then dehydrated with series of graded ethanol concentration (95%, 95%, 100%, and 100%) and slide mounted. Color of the antibody staining in the tissue sections were observed under 40x magnification in a compound microscope (Axiovert, Carl Zeiss, USA). The staining intensities were quantified using IHC Profiler, a plugin in ImageJ software, to determine the H-score (Histo score) that is directly proportional to the concentration of DAB, as described by Varghese F, et al (Varghese et al., 2014).

### 2.10 Primary culture of astrocytes

Primary astrocytes were isolated from the collected brain tissues as described previously (Schildge et al., 2013). Briefly, the brain tissues of all the experimental groups were isolated, immediately after decapitation. Once the blood vessels and meninges were removed, the forebrain tissues were finely chopped in the culture medium, EMEM. Subsequently, the minced tissue was dissociated using 0.05% trypsin at 37^0^C for 30 min. Trypsinization was stopped by adding 10% (v/v) FBS in EMEM. To isolate individual astrocytes, the cell suspension was passed through 40 μm strainer. At last, astrocytes were seeded at a density of 1.5×10^5^ cells/cm^2^ in 90% EMEM containing 10% FBS, and 4.5 g/L glucose. The astrocytes were cultured at 37°C with 95% air and 5% CO_2_.

### 2.11 Measurement of ROS

Intracellular generation of ROS regulated by the drugs, either alone or in combination on glioma cells, was measured by calculating the fluorescence intensity of 2’,7’ -dichlorofluorescein diacetate (DCF-DA) oxidized product. Briefly, 1×10^3^ astrocytes isolated from each experimental group were seeded on a 96-well plate. The cells were then stained with 100 μl of DCF-DA (100 μM) and incubated at 37^0^C for 15 min. The relative fluorescence intensity of oxidized product of DCF-DA was measured by reading the absorbance at 530 nm in a spectrophotometer (Molecular Devices Spectra-Max M5, USA).

### 2.12 Measurement of antioxidant, non-antioxidant enzymes and lipid peroxidation

Biochemical analysis were performed on the brain tissue lysates to measure (i) Superoxide dismutase (SOD) activity using SOD Assay kit (Cat No. 706002); (ii) Catalase activity using CAT Assay Kit (Cat No.707002); (iii) Glutathione Peroxidase (GPx) activity by GPx Assay Kit (Cat No. 703102); (iv) Glutathione activity with GSH Assay Kit (Cat No.703002) and for the measurement of Malondialdehyde (MDA) levels, which is a by-product of lipid peroxidation, the Thiobarbituric acid reactive substances (TBARS) assay Kit (Cat No. 10009055) was employed. All the biochemical parameters were performed using the kits available from Cayman chemical, Ann Arbor, USA. All the analyses were performed according to the manufacturer’s protocol. Three independent biological replicates were performed for each assay.

### 2.13 Quantitative real-time polymerase chain reaction (q-RT PCR)

The total RNA from the brain tissues of all experimental groups were isolated using TRIZOL reagent (Takara Bio, Shiga, Japan) and the respective complementary deoxyribonucleic acid (c-DNA) was synthesized using Hi-c-DNA Synthesis Kit (HiMedia, India). Quantitative RT-PCR was carried out using a CFX96 thermo cycler (Bio-Rad, California, USA) and TB Green Premix Ex Taq I (Takara Bio, Shiga, Japan) to detect messenger ribonucleic acid (mRNA). The specific PCR primer sequences used in this study were designed using Basic Local Alignment Search Tool (BLAST) (The primer sequences used are listed in **supplementary Table S1**). Independent experiments were conducted in triplicate. The cycle threshold (Ct), representing a positive PCR result, is defined as the cycle number at which a sample’s fluorescence intensity crossed the threshold automatically determined by the CFX96 (Bio-Rad, California, USA). The relative changes in gene expression were calculated with the 2^-ΔΔCt^ method.

### 2.14 Cell lysis and protein amount quantification

For protein isolation, 100 mg of brain tissues of all the experimental groups were washed with 0.9% NaCl, 5 times each, to remove contaminated blood and was fully grilled in liquid nitrogen. 2 ml of RIPA protein extraction buffer (20 mM Tris-HCl (pH 7.5), 150 mM NaCl, 1 mM sodium ethylenediamine tetra acetate (EDTA), 1 mM EDTA, 1% NP-40, 1% sodium deoxycholate, 2.5 mM sodium pyrophosphate, 1 mM β-glycerophosphate, 1 mM Na3VO4, 1 μg/ml leupeptin) was added to the brain tissues, crushed and then centrifuged at 12000 ×g for 15 minutes at 4°C. The lysates were then collected, centrifuged again at 12000 ×g for 15 minutes at 4°C. The protein extract collected as a supernatant was then quantified using Bradford method (Bradford, 1976).

### 2.15 Western Blotting

Following quantification, 40-100 μg of the total protein were separated by sodium dodecyl sulphate polyacrylamide gel electrophoresis (SDS-PAGE) and then transferred onto polyvinylidene difluoride (PVDF) membrane (Millipore, Germany) by the wet transfer method. The membrane was blocked with 5% skim milk at room temperature and then incubated with the indicated primary antibodies against β-actin, Nrf2, hypoxia inducible factor-1 alpha (HIF-1α), PI3K, pyruvate dehydrogenase kinase-1 (PDK1), pAKT1, phosphatase and tensin homolog (PTEN), Caspase-9, pmTOR and BAD (Elabsciences, Wuhan, China) on an orbital shaker at 4°C overnight. After washing three times with 1X-tris-buffered saline/tween-20 (TBST), the blots were incubated with HRP-conjugated secondary antibody (Elabsciences, Wuhan, China) for 1 hour on an orbital shaker at room temperature. Each immune complex was detected by Enhanced Chemiluminescence (ECL) Substrate (Santa Cruz, Texas, USA) and was visualized by the Chem-doc (Pearl® Trilogy/ LI-COR, USA). Quantitative analysis of Western blot band intensities was made by ImageJ® software.

### 2.16 Enzyme-Linked Immune Sorbent Assay

According to the manufacturer’s protocol, for the isolated protein samples from the tissue homogenates of all the experimental groups, Enzyme-Linked Immune Sorbent Assay (ELISA) was performed for (i) VEGF (Cat No. E-EL-R1058); (ii) VEGFR-1 (Cat No. E-EL-R0911); (iii) Caspase-8 (Cat No. E-EL-R0280) and (iv) Caspase-3 (Cat No. E-EL-R0160). All the ELISA kits were procured from Elabsciences, Wuhan, China. The concentration levels of the above-mentioned proteins were calculated from a standard curve obtained.

### 2.17 Cell Cycle analysis

For cell cycle analysis, 1×10^6^ isolated astrocytes from the brain tissues of each experimental groups were rinsed with ice-cold PBS and re-suspended in 100% methanol at 4°C for 40 minutes. The astrocytes were then pelleted via centrifugation at 2000 rpm for 5 minutes, re-suspended in 500 μl of PBS containing 0.1% triton X-100 and 22 μg of 4’,6-diamidino-2-phenylindole (DAPI), and incubated in dark for 30 minutes at 25^0^C. Astrocytes were fully re-suspended and the cell cycle pattern was analyzed using BD Celesta Flow Cytometry (Becton-Dickinson, California, USA). The results were further analyzed and quantified using FlowJo 7.6 software (Beckman Coulter, CA, USA).

### 2.18 Annexin V/ 7’AAD assay

Percentage of apoptotic cells were determined by flow cytometry to measure the apoptotic rate after staining with Annexin V and 7-amino-actinomycin D (7’AAD) kit (Annexin V-FITC - AAD Kit, Becton-Dickinson, USA). Briefly, 1×10^6^ astrocytes isolated from all the experimental groups were harvested and washed in ice-cold PBS twice. The astrocyte pellets were resuspended in Annexin V binding buffer and incubated on ice for 5 minutes. After which, the cells were stained with 7’AAD and incubated for 5 minutes, according to the manufacturer’s protocol. The rate of cell apoptosis was performed by Cyto-FLEX S Flow Cytometer and was analyzed using FlowJo 7.6 software (Beckman Coulter, CA, USA).

### 2.19 Molecular docking analysis

To perform molecular docking studies, an evaluation version of Schrödinger software package was used (Schrödinger Release 2020-3: Maestro, Schrödinger, LLC, New York, NY, 2020). Crystal structure of human VEGFR1 (3HNG) having a resolution of 2.7 Å was retrieved from protein data bank (Berman et al., 2002). Prior initiating the docking protocol, protein structure was minimized using the Protein Preparation Wizard with optimal potential to create the liquid simulation (OPLS)-2005 force field (FF). Also potentially occurring stereochemical short contacts in the protein structure were removed during the protein preparation. Subsequently water molecules without any contact were removed followed by addition of hydrogen atoms to the structure, principally at the sites of the hydroxyl and thiol hydrogen atoms, to correct ionization as well as tautomeric states of the amino acid residues. The 2D structure of drug molecules T (Pub-chem ID: 5394) and its metabolites 3-methyl-(triazen-1-yl) imidazole-4-carboxamide (MTIC) (Pub-chem ID: 54422836) and 5-amino-imidazole-4-carboxamide (AIC) (Pub-chem ID: 9679), M (Pub-chem ID: 14219) and its metabolite Guanylurea (GUA) (Pub-chem ID: 8859) and E (Pub-chem ID: 65064) were downloaded from PubChem in .mol file and prepared using the ligprep module (**Figure 5**). Prior performing docking protocol using Glide standard precision (SP), the binding site pocket and grid for docking were generated using the ligand bound to the native X-ray crystallographic structure of 3HNG.

### 2.20 Density-Functional Theory (DFT) Calculations

The best binding poses of the six molecules docked with 3HNG, were geometry optimised using the Jaguar panel and subjected to DFT analysis using the Becke’s three-parameter exchange potential and Lee-Yang-Parr correlation function (B3LYP) theory with 6-31G** level as the basis set (Gill et al., 1992; Selvaraj et al., 2021a). This helps to compute the reactivity and stability of the molecules (Zheng et al., 2013).

### 2.21 Statistical analysis

Data were analyzed using GraphPad Prism 8 (San Diego, USA). Statistical analysis was carried out using by Student’s t-test or one-way/ two-way ANOVA followed by Dunnet’s/ Wilcoxon-Gehan multiple comparing tests.

## 3. Results

### 3.1 Triple-drug combination therapy depicts superior survival benefit without any organ-based toxicity

To study the beneficial mechanism of these three different drugs, both individually and in combination, in terms of prolonging prognosis and survival rate, a survival study was carried out. The animal-grouping used in this study is depicted in **Figure 1A (* Survival study; # Experimental studies)**. An orthotopic glioma model was developed as explained in methods section. 20 days after tumor implantation, the rats were treated with the indicated dose of drugs, both individually and in combination. Subsequently, the effect of drugs on prolonging the prognosis of TC and treated rats, was followed up for 25 weeks as reported previously by Tseng et al (Tseng et al., 2016).

At the end of the study, the rats were sacrificed and the survival results were interpolated using Kaplan-Meier Survival graph. As shown in **Figure 1B**, treatment of the rats with the triple-drug combination (TME) significantly improved tumor suppression and prolonged the duration of survival (>25 weeks) in 50% of the treated animals, relative to the effect observed with the other experimental groups. The median survival rate of the triple-drug treated animals (TME) (24 weeks) was significantly higher than the tumor-control group (7.5 weeks) (P<0.0001). Further, among the dual-drugs administered, TM and TE showed a slightly better median survival rate of 18.5 and 15 weeks respectively relative to ME group (13.5 weeks). Conversely, delivery of either of these drugs as individual regime did not correlate with significant improvement in the survival of rats when compared with the triple-drug combination. These results demonstrated that combined delivery of T, M and E produced superior survival benefit and prolonged the survival of animals.

Following glioma implantation and successive treatment of the glioma-induced rats with the drugs, both individually and in combination, histopathological analysis of vital organs was carried out by H&E staining. Interestingly, no significant pathological changes were observed in any of the organs obtained from the drug-administered groups when compared with the control group. This result highlights that the none of the treatment strategies induced any organ-based toxic effects (**Supplementary Figure S1**).

### 3.2 Triple-drug combination reduced the number of tumor cells

Following assessment of the survival benefit bestowed within the combinatorial treatment, we were interested in determining their effect on pathological changes in the brain tissues.

Interestingly, histological analysis by H & E staining depicted a considerable decrease in tumor cells in the sections obtained from the triple-drug combination (TME) when compared with the other treatment groups and control. While the tumor-control (TC) rats depicted intense cellularity and extensive haemorrhage in blood vessels (**Figure 1C-b**), the cellularity and haemorrhage were drastically reduced in the rats treated with the triple-drug combination (**Figure 1C-i**). In fact, the tumor cell density in the triple-drug treated tissues were almost similar to the non-tumor brain regions. Among the dual-drug treatment groups, TM and TE portrayed a slight reduction in the whorled structures and reduced cell density but were not as effective as the triple-drug combination (**Figures 1C-f & g**). Though treatment with these drugs as individual regime had relatively modest effect with decreased haemorrhage, they were not as effective as the triple-drug combination. These results shed light on the fact that besides prolonging survival, the triple-drug combination can effectively reduce the extent of tumor, which was grossly observed in a significant manner.

### 3.3 Triple-drug combination reduced proliferation and angiogenesis

Considering the significant anti-invasive effect of the triple-drug combination, we then evaluated the expression levels of VEGF (angiogenic marker) and Ki67 (proliferation marker) in the tumor sections of the experimental groups. Assessment of cytoplasmic positivity for VEGF was significantly reduced in tumors treated with the triple-drug combination (6.36%) (specified by red arrows in **Figure 2A-i**), when compared with tumor-control group, which exhibited the highest (70.61%) level of cytoplasmic positivity (black arrows indicated in **Figure 2A-b**). Hitherto, TM, T-alone and TE depicted a moderate cytoplasmic positivity, as evident by the H-score of, 38.59, 50.38 and 46.45% respectively (**Figure 2A-j**).

**Figure 2.**
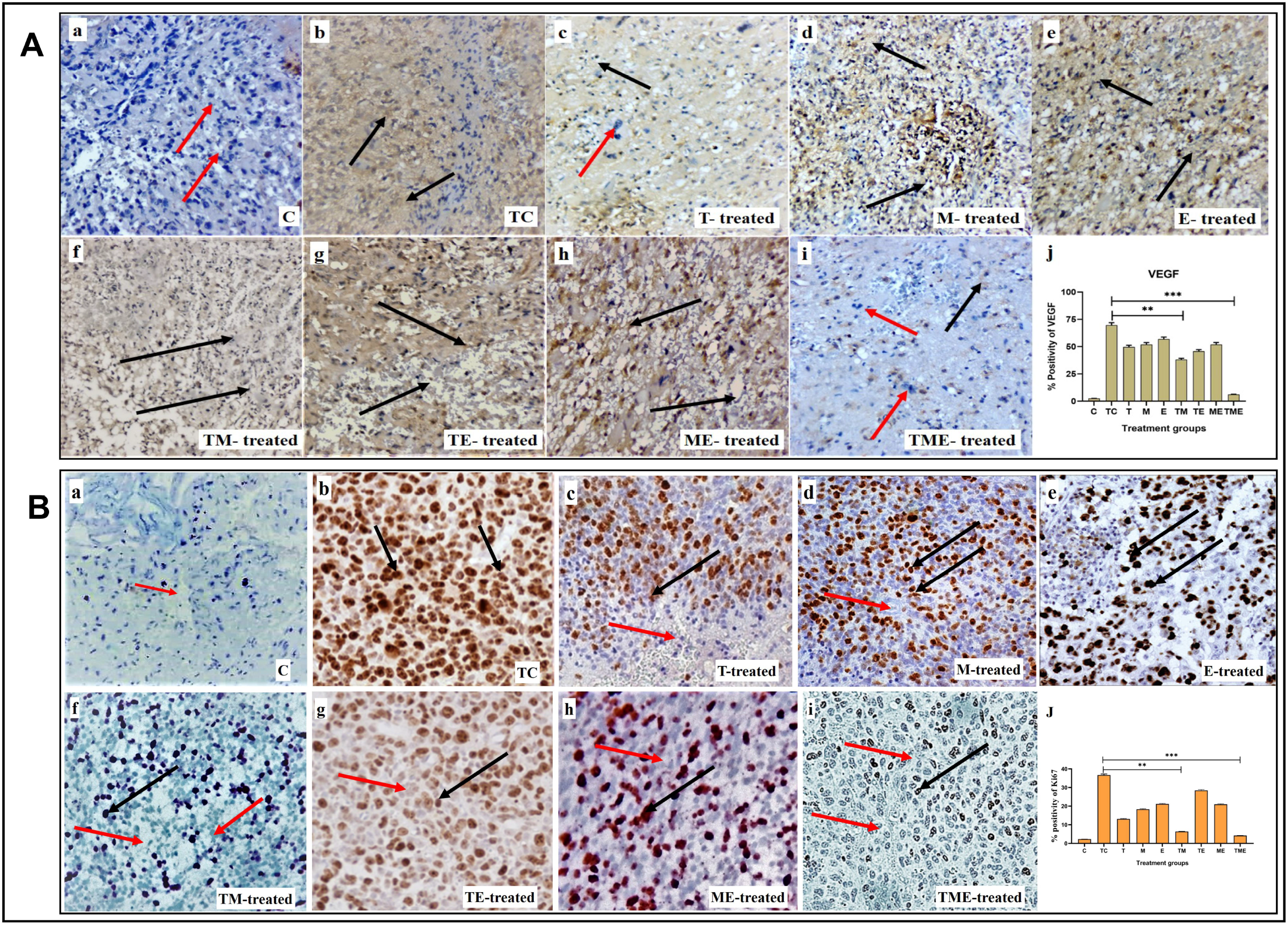
(**A**) IHC staining for VEGF in tumor sections from different experimental groups at 40x magnification. (A**-a** to **A-i**) IHC staining for VEGF in various treatment groups, wherein cytoplasmic positivity is denoted by black arrows and cytoplasmic negativity is denoted by red arrows respectively. (**A-j**) Quantification of cytoplasmic positivity for VEGF in each group. (**B**) Ki67 nuclear expression in different experimental groups at 40x magnification. (**B-a** to **B-i**) IHC staining for Ki67 in various treatment groups, wherein nuclear positivity is denoted by black arrows and nuclear negativity is denoted by red arrows respectively (**B-j**) Quantification of nuclear positivity for Ki67 in each group (**P<0.001, *** P<0.0001).3.4. Triple-drug combination significantly enhanced the ROS levels and levels of antioxidant and non-antioxidant enzymes in glioma.

In addition, Ki67 exhibited a lower nuclear positivity rate in the tumor-induced rats that received the triple-drug combination therapy (4.21%) (as specified by red arrows in **Figure 2B-i**). Whereas, the highest nuclear positivity rate was noticed in tumor-control rats (36.73%) (as indicated by black arrows in **Figure 2B-b**). However, the dual-combination therapy of TM (6.42%), T-alone (13.14%) and M-alone (18.36%) disclosed significant lower nuclear positivity of Ki-67 expression, but were not as significant as triple-drug combination (**Figure 2B-j**). Conversely, the other treatment groups did not induce any significant changes in the nuclear positivity of Ki-67 expression (P<0.01).

In most cancers, angiogenesis is vital for tumor growth and proliferation, which is often facilitated by reduced levels of ROS, while conversely elevated ROS levels have been reported to reduce tumor progression and inhibit angiogenesis (Kumari et al., 2018). Therefore, to assess whether the drugs induced oxidative stress, we measured the intracellular ROS generation by measuring the DCF fluorescence intensity. This method is commonly used in ROS investigations and is based on the application of H2DCFDA (acetylated form of DCF), which is consecutively deacetylated inside the cells by intracellular esterase. The resulting molecule is oxidized by intracellular ROS to produce a fluorescent product, DCF. Treatment of glioma-induced rats with the triple-drug combination were associated with a significant increase in ROS (P<0.001) as compared to that of tumor-control rats (**Figure 3A**). Further, the dual-drug combination (TE) was also shown to exhibit a profound increase in the ROS levels (P<0.01) but were not as significant to triple-drug combination (**Figure 3A**). Though the other treatment groups exhibited an increase in the intracellular ROS levels, yet they were not significant (**Figure 3A**). These results demonstrated that the triple-drug combination treatment triggered ROS production in glioma-induced rats.

**Figure 3.**
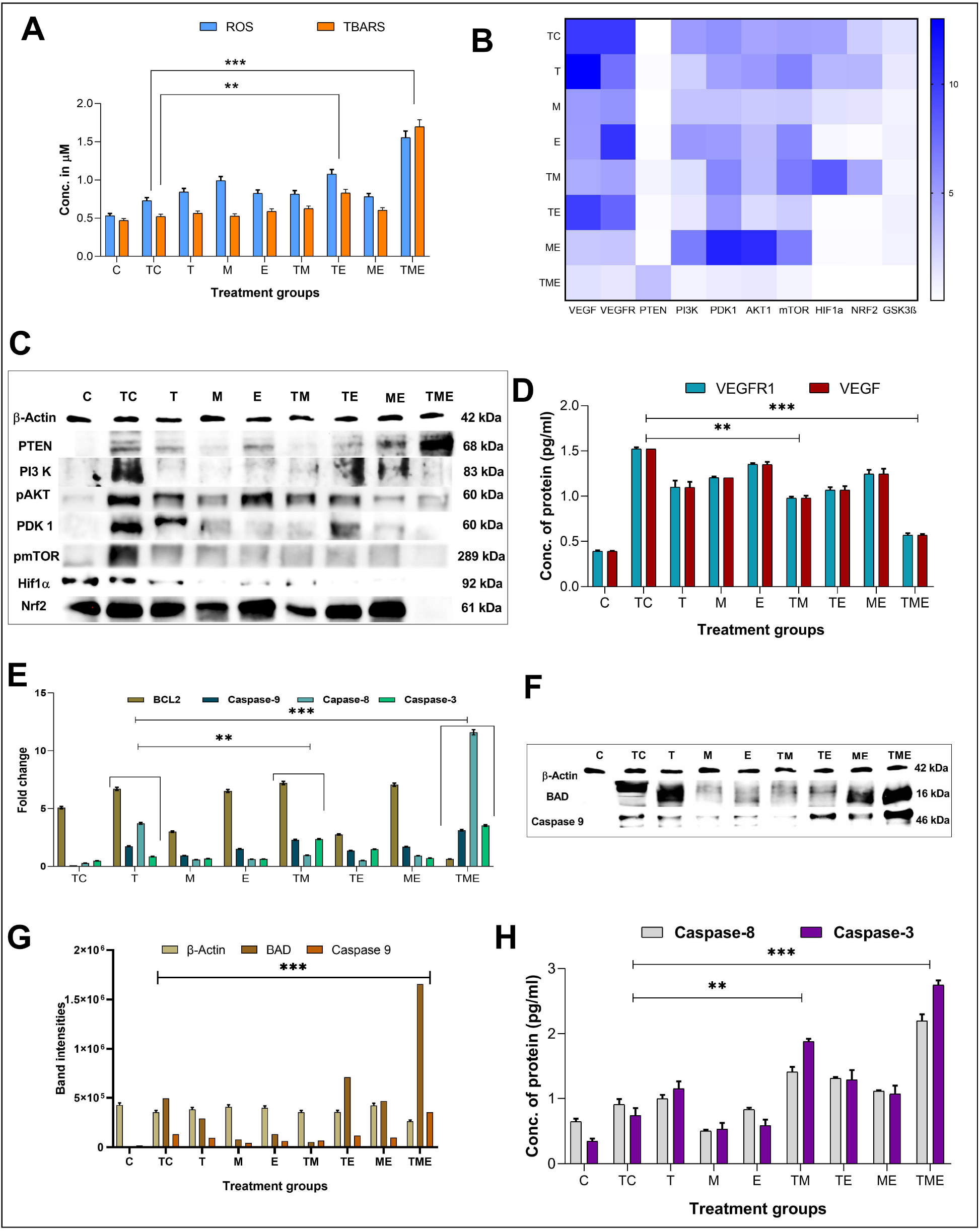
(**A**) shows the cumulative ROS and TBARS activity observed in different treatment groups, in which the triple-drug treatment and the dual-drug treatment of TE significantly elevated both ROS and TBARS activity. (**B**) Heat map for the gene expression levels of VEGF, VEGFR, PTEN, PI3K, PDK1, AKT1, mTOR, GSK3β PTEN, HIF-1α and Nrf2 the significance was calculated using two-way ANOVA. (**C**) Western blot bands for markers of PI3K/AKT/GSK3β/Nrf2 pathway (HIF-1α, Nrf2 PTEN, PI3K, pAKT1, PDK1 and pmTOR) analysed for various treatment groups. (**D**) Protein levels of VEGFR1 and VEGF by ELISA. (**E**) Gene expression levels of apoptotic markers (BCL2 and caspase-9, 8 & 3). (**F**) Western blot bands for the apoptotic proteins (BAD & caspase-9) proteins analysed for various treatment groups. (**G**) The intensities of apoptotic proteins BAD and caspase-9 band were calculated against β-Actin. (**H**) Protein levels of caspase-8 & 3 were analysed using ELISA. The significance was calculated using two-way ANOVA (**P<0.001, *** P<0.0001).

ROS formed in tumor cells results in lipid peroxidation and subsequently increases MDA. MDA can be directly quantified by determining the levels of TBARS, a marker of lipid peroxidation. **Figure 3A** depicts the concentration of TBARS in the brain tissues of all the experimental groups. In the present study, the levels of TBARS were significantly enhanced in the triple-drug combination (P<0.001) and the dual-drug combination of TE (P<0.01) (**Figure 3A**) when compared to tumor-control rats. However, the levels of TBARS were not that significant as of triple-drug combination than the other experimental groups.

On the other hand, we were also interested in determining the effect of the chosen drugs both individually and in combination on the major antioxidant and non-antioxidant enzymes like SOD, CAT, GPx and GSH, as enhancement of these antioxidant enzymes indicates an increased accumulation of super oxides, such as O_2_^-^, H_2_O^-^ etc., leading to enhanced oxidative stress. Interestingly, we observed that the activity of all the antioxidant and non-antioxidant enzymes were significantly enhanced in the triple-drug combination (P<0.001) treated rats when compared with that of tumor-control rats. However, a similar effect was observed in the dual-drug combination (TE) and E-alone (P<0.01) treated rats, but were not as significant as triple-drug combination treated rats (**Supplementary Figure S2**). Since glioma is often associated with superoxide and peroxides mediated chemo-resistance to T, co-administration of M and E had significantly enhanced the susceptibility of glioma cells to T via elevating the ROS levels, enhancing lipid peroxidation and antioxidant potential.

### 3.4 Triple-drug combination effectively reduced antioxidant defence

The expression of some of the antioxidant defence genes mentioned above are regulated by the transcription factor Nrf2 under hypoxic conditions, allowing cells to regulate the oxidative stress-mediated ROS species. This is brought about by binding of Nrf2 to the promoter regions of the antioxidant response elements, which in turn reduces the levels of ROS in the cells. The formation and accumulation of ROS in hypoxic cells is a hallmark of hypoxia involved in blood brain barrier (BBB) dysfunction. In association with the production of ROS, several factors associated with hypoxia are also activated or stabilized. Effectively, HIF-1, more specifically HIF-1α, is stabilized and stimulates the transcription of genes involved in various processes such as angiogenesis, cell proliferation, inflammation or cancer. Owing to the fact that both Nrf2 and HIF-1α, are well established as mechanisms of resistance to anticancer therapies, simultaneously targeting these pathways would represent an attractive approach for therapeutic development. Interestingly, in the present study, we observed that the triple-drug combination significantly attenuated the gene expression levels of Nrf2 and HIF-1α, thereby deactivating the antioxidant defence mechanism in tumor cells (P<0.0001). In addition, treatment with dual-drug combination, TE and E-alone also exhibited reduced expression of these genes, but were not as potent as triple-drug combination (**Figure 3B**). These results put forth the fact that, the toxicity effect of this triple-drug combination is directly related to their ability to inhibit antioxidant defence and trigger apoptosis via inhibition of Nrf2 and HIF-1α.

### 3.5 Triple-drug combination inhibits the PI3K/AKT/GSK3β/Nrf2 pathway

Since there was reduction in the gene expression levels of Nrf2 and HIF-1α by the triple-drug combination, we were further interested on focusing in the molecular mechanism involving the upstream signalling pathways of Nrf2 and HIF-1α. Nrf2 and HIF-1α are the main targets for VEGF/PI3K/AKT and GSK3β, wherein p-AKT and GSK3β promotes the separation of Nrf2 from Keap1, thereby leading to the translocation of Nrf2 into the nucleus. GSK3β is a substrate of the PI3K pathway that is constitutively active in unstimulated cells and is known to participate in the protective cellular response to oxidative stress. Thus, VEGF/PI3K/AKT/GSK3β signalling pathway may be one of the key regulators of cell survival. PTEN, which is the downstream molecule of this pathway, acts as a tumor suppressor by inhibiting tumor cell growth and enhancing cellular sensitivity to apoptosis. Loss of PTEN activity leads to the permanent PI3K/AKT pathway activation. As shown in **Figure 3B**, the triple-drug combination significantly (P<0.0001) reduced the gene expression levels of VEGF, VEGFR, PI3K, PDK1, AKT1, mTOR and GSK3β while it significantly (P<0.0001) enhanced the expression of PTEN, when compared with the tumor-control and other treatment groups. Similarly, the dual-drug combination, TM and TE significantly (P<0.001) reduced the levels of VEGF, VEGFR, PI3K, PDK1, AKT1 and mTOR, but not to the extent of triple-drug combination. Remarkably, these dual-drug combinations mentioned did not have any significant effect on the levels of PTEN. However, the other treatment groups did not show any significant changes on the gene expression levels of the PI3K signalling pathway when compared to the tumor-control (**Figure 3B**). From this, it was evident that our triple-drug combination significantly reduced glioma proliferation by inhibiting the Nrf2/HIF-1α-mediated PI3K/ GSK3β signalling pathway.

### 3.6 Triple-drug combination synergistically modulated the activation of PI3K/AKT/ GSK3β/Nrf2 pathway in glioma-induced rats

In order to elucidate if the results observed from our gene expression studies reconciled at the protein expression levels as well, the protein levels of VEGF/PI3K/AKT/GSK3β/Nrf2/HIF-1α pathway were assessed by immunoblotting and ELISA. The results revealed that the triple-drug combination markedly inhibited the protein expression of VEGF, VEGFR, PI3K, p-AKT, PDK, p-mTOR, Nrf2 and HIF-1α when compared to that of tumor-control (P<0.001) (**Figure 3C**); while it significantly enhanced the expression of PTEN. This was in concordance with the gene expression analysis. Likewise, the dual-drug combination (TM and TE) too significantly decreased the levels of VEGF and VEGFR (P<0.01) (**Figure 3D**), but was not as effective as triple-drug combination. Besides, the dual-drug treatment (TM) reduced the protein expression of PI3K (P<0.01), while the levels of p-AKT1 were significantly reduced by the dual-drug combination of TM and TE as well as in the individual treatment of M. In addition to the triple-drug combination, the individual treatment with M significantly reduced the levels of p-mTOR, indicating a moderated cell growth and proliferation. However, the dual-drug combination (TM, TE and ME) significantly reduced the levels of Nrf2, conversely, they did not have any significant effect on the levels of HIF-1α (P<0.01). Surprisingly, individual treatment with E significantly reduced the levels of both Nrf2 and HIF-1α (P<0.01) (**Figure 3C).** These results demonstrated that the triple-drug combination inhibited glioma cell growth via VEGF/PI3K/AKT/GSK3β/Nrf2 signalling pathway.

### 3.7 Triple-drug combination induced apoptosis in glioma cells

Previously, it has been reported that increased oxidative stress and reduced proliferation via inactivation of PI3K/AKT/mTOR pathway is often linked to the induction of apoptosis in glioma (Li et al., 2016). Hence, to gain deep insights on the effect of our drugs on induction of apoptosis, following their efficacy on oxidative stress and proliferation, we analysed the gene and protein expression of the mediators of the intrinsic pathway of apoptosis namely, BAX, BAD, caspase -9, 8 and the executioner caspase-3 in the tumor tissues of the treated rats. As expected, our gene expression results demonstrated that the triple-drug combination (TME) significantly reduced the levels of BCL2 and enhanced the expression levels of BAX, BAD, caspase-9, 8 and 3 when compared to that of tumor-control group (P<0.0001). The dual-drug combination of TM treatment significantly (P<0.001) reduced the levels of BCL2 and enhanced the levels of BAX, BAD, caspases-9, 8 & 3 (P<0.001) (**Figures 3E-H, Supplementary Figure S3)**. A similar trend in the expression levels of apoptotic proteins was observed by Western blot and ELISA, however, from our western blot results it was noted that T had a significant role in the elevating the levels of BAD as BAD levels in both T and TME groups were almost equally higher. From this, it was evident that the triple-drug combination of TME, induced apoptosis by activating the BAD/BAX and caspase complex, thereby suggesting a reduced proliferation and induction of apoptosis at gene and protein level.

### 3.8 Triple-drug combination alters the cell cycle distribution pattern in glioma tissues

To further evaluate if the inhibition of glioma proliferation and apoptosis was accompanied by alterations in the cell cycle pattern, flow-cytometry was performed in the cells isolated from glioma-induced rats. Interestingly, we found that when compared to tumor-control, the triple-drug combination significantly enhanced the fraction of non-proliferating cells (G1 phase) by 21.5% (62.3% vs. 83.8%) while significantly decreasing the count in proliferating cells (G2/M phase) by 8% (**Figures 4A-D**); Also, the triple-drug combination significantly reduced the number of cells in S phase by 13.5% (29.1% vs. 15.6%) when compared to the tumor-control group (P<0.0001; **Figure 4D**). Alongside, the individual treatment with TM depicted a significant increase in the number of cells in G1 phase by 20.4 & 19.4% respectively, however not much effect was observed on the proportion of the cells in S-phase when compared to that of TC group (P<0.0001). From our cell cycle analysis, it was clear that, besides inducing oxidative stress-mediated inactivation of PI3K/AKT/mTOR pathway, the combinatorial treatment significantly arrested the cells at G1 phase, thus leading to apoptosis (**Figures 4A-D**).

**Figure 4.**
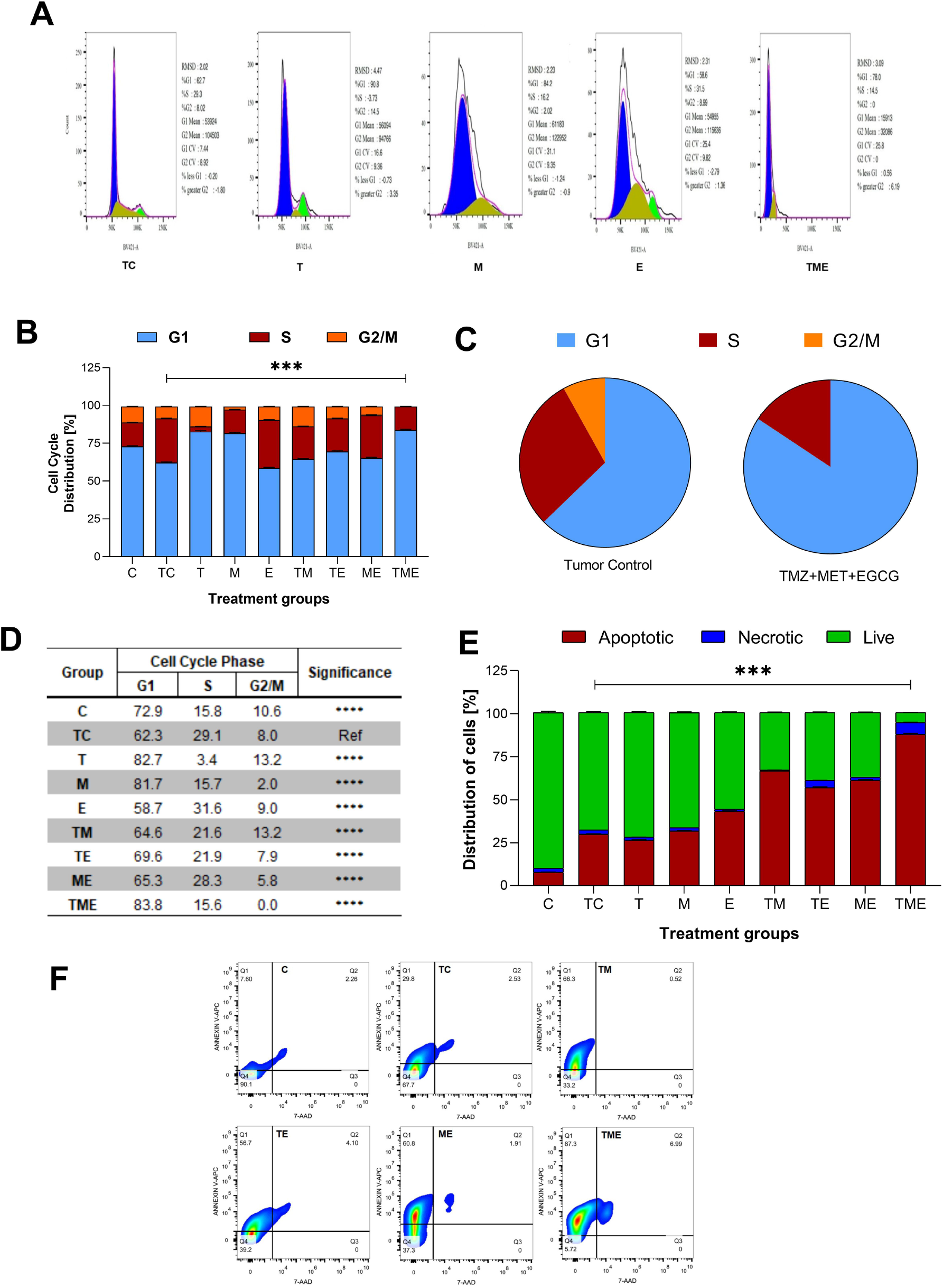
Flow Cytometric analysis. (**A**) Effect of the drugs, individually and in combination on cell cycle, wherein the histograms represent the DNA content of the cells in each of the cell cycle phases (G0/G1, S and G2/M). Different cell cycle phases were plotted (**B**) Percentage of total events, wherein the bar-graph depicting the percentage of cells in G1, S and G2/M phases in each treatment group. (**C**) Pie chart representing the percentage of cells in G1, S and G2/M phases in tumor-control and triple-drug treatment group respectively. (**D**) Tabulation of percentage of cell in different phases in each treatment group. (**E**) The clustered bars represent total percentages of live, apoptotic and necrotic cells for the various treatment groups. (**F**) Represent the cell population in the 4 different quadrants (Q1- early apoptotic, Q2- necrotic, Q3- late apoptotic and Q4- live cells).

### 3.9 Triple-drug combination increased the number of apoptotic cells in glioma tissues

Following the oxidative stress-mediated inactivation of PI3K/AKT/mTOR pathway leading to apoptosis at molecular level, we were further interested to determine the underlying mechanism of cell death induced by our drugs both individually and in combination. Hence, the astrocytes isolated from various treatment groups were subjected to flow cytometric analysis by Annexin V/ 7’AAD staining. It is well-known that annexin V binds to early apoptotic cells (Q1), whereas 7’AAD binds to late apoptotic cells (Q3); Further, the population of cells which contain both annexin V and 7’AAD are considered to be necrotic ones (Q2) respectively. Live cells are found in Q4. Given this fact, our results highlighted that compared to TC, the triple-drug combination significantly enhanced the number of apoptotic (30% vs. 87%) and necrotic cells (2.5% vs. 7%) by 57 % (Q1) & 4.5% (Q2) respectively and significantly reduced the number of live cells (68% vs. 6%; Q4) (P<0.001) (**Figures 4E and 4F**). Similarly, the dual-drug combination of TM too enhanced the levels of apoptotic cells by 36.5% and significantly reduced the number of live cells by 34.5% as compared to the tumor-control group (P<0.01; **Figure 4F**). This undoubtedly confirmed that the triple-drug combination promoted cellular death in glioma via apoptosis.

### 3.10 Molecular docking

The protein structure after minimization was found to be satisfactory to proceed further with the docking analysis **(Figure 5A)**. The outcome of VEGFR1 interaction with selected drug molecules was evaluated using the Glide score. **Figure 5B** shows the interaction profile of selected drug molecules with VEGFR1. Based on the glide scores of the interaction of the six different molecules 3HNG, T ranked top (glide score, -7.938 kcal/mol), with two hydrogen interactions, one between Cys 912 and oxygen atom at the fourth position of ligand’s tetrazine ring and second between Asp 1040 and NH2 at the eighth position of ligand’s imidazole ring with a bond length of 1.68 and 2.28 Å, respectively. While MTIC ranked second (−6.829 kcal/mol), demonstrating two hydrogen bond interactions, one between Cys 912 and ligand’s carboximide group and second between Asp 1040 and ligand’s aminodiazenyl moiety with a bond length of 1.8 and 2.11 Å, respectively. Furthermore, a π-π interaction was noted between the benzene ring of Phe 1041 and ligand’s imidazole ring with a bond length of 4.72 Å. AIC was ranked third (−5.736 kcal/mol), and exhibited two hydrogen bond interactions, one between Val 907, and NH of carboxamide and second hydrogen bond between Glu 878 and NH of imidazole ring with a bong length of 2.53 and 1.9 Å, respectively. Also, an π-cation interaction between Lys 861 and ligand’s imidazole ring was noted with a bond length of 4.32 Å. Whereas, E demonstrated three hydrogen bond interaction, one each between the hydroxyl group and carbonyl group of ligands trihydroxybenzoate moiety with Asp 1040 and Arg 1021 of 3HNG with a bond length of 2.19 and 1.81 Å, respectively. Third hydrogen bond interaction was noted between the hydroxyl group of trihydroxy-phenyl attached to di-hydrochromenyl moiety and Asp 807 with a bond length of 2.0 Å. In addition, an π -π interaction was observed between the benzene ring of dihydrochromenyl moiety and the imidazole ring of His 1020 with a bond length of 5.44 Å. Compound GAU demonstrated three hydrogen interactions of which two interactions were noted between the ligand’s NH_2_ group of carbamoyl-amino portion, and the active site residues namely His 1020, and Asp 1040 with a bond length of 1.75 and 1.63 Å, respectively. The third bond exhibited between ligand’s NH group and Asp 1040 with a bond length of 2.27 Å. Finally, of the six compounds investigated for the interaction, it was noted that M revealed two hydrogen bond formation one between Glu 878 and second between Asp 1040 with a bond length of 1.85 and 2.28 Å, respectively. The details of all the drugs along with their active metabolites and their respective glide scores are given in **Table 1**.

**Figure 5.**
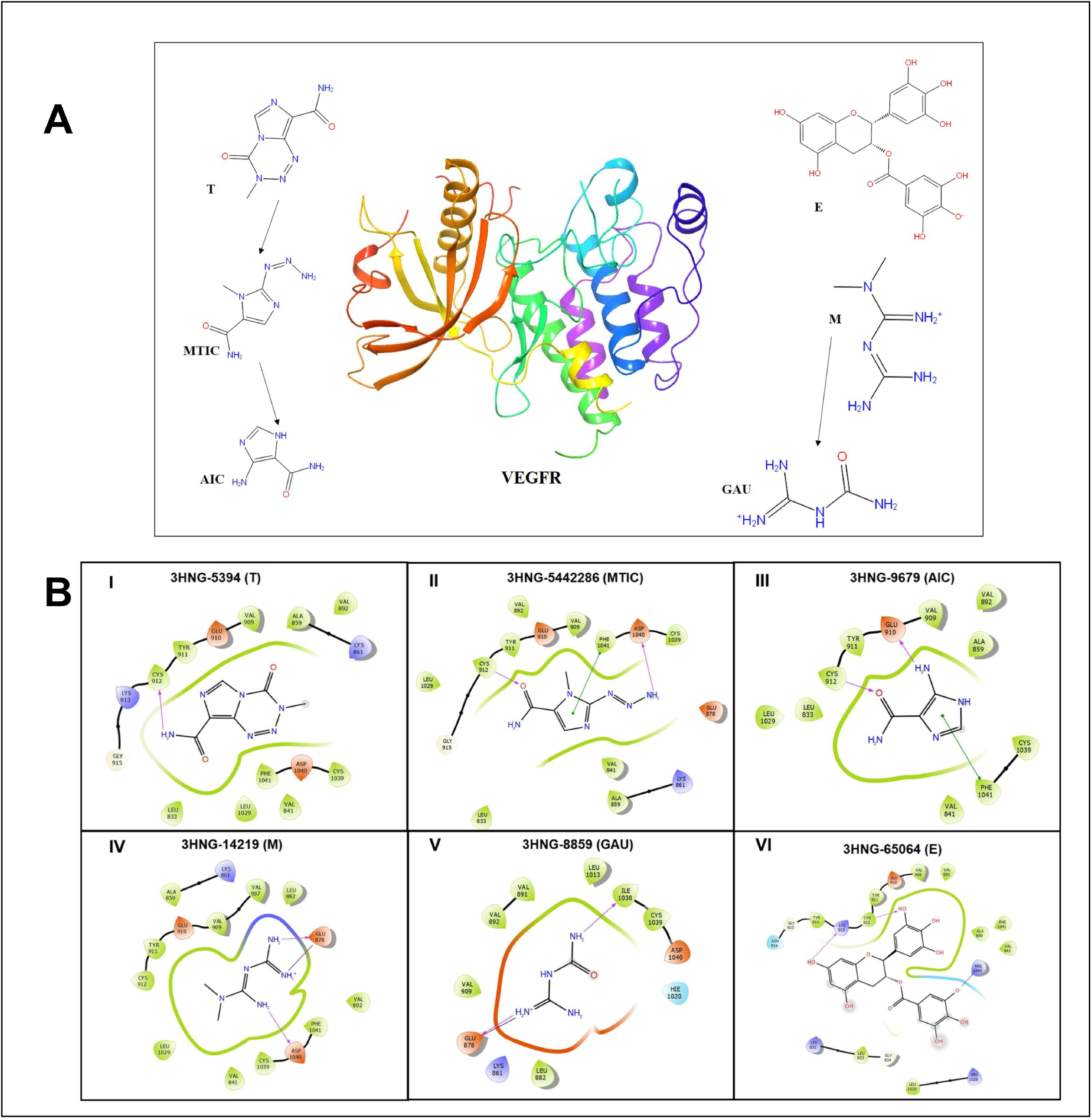
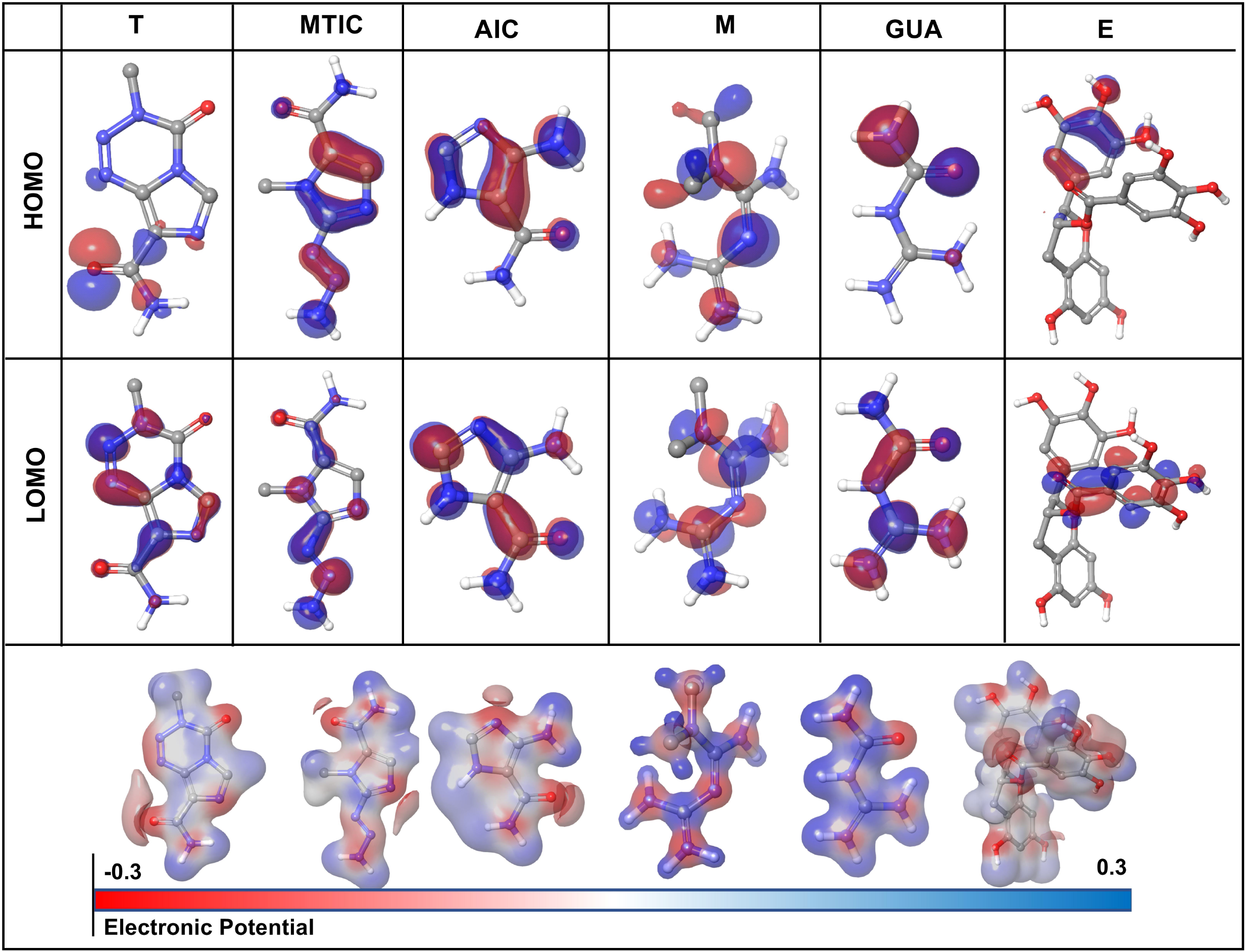
(A) 3D structure of human VEGFR protein (3HNG) and 2D structures of the drugs employed in docking. (B) Site maps of drug-protein reactions in the VEGFR active domains. The drugs are arranged in ascending order of their glide score where, (I) T, (II) MTIC, (III) AIC, (IV) M (V) GUA and (VI) E.

### 3.11 DFT Calculations

The HOMO (highest occupied molecular orbitals) and LUMO (lowest unoccupied molecular orbital) energies and the corresponding energy gap of the drugs and their metabolites namely T, MTIC, AIC, M, GUA and E are listed in **Table 3** and **Figure 6** along with the molecular electrostatic potential map. This energy gap indicates the electronic excitation energy and has a significant role in the stabilisation of interaction between the receptor protein and the drug molecule (Haas et al., 2018; Zheng et al., 2013). The red colour regions in the plots indicate the electropositive regions while the blue regions indicate the electronegative regions.

## 4. Discussion

Glioma, arising from a multistep tumorigenesis of the glial cells is usually characterized by dismal prognosis (Li et al., 2016). Despite decades of research, the overall survival in most of the glioma cases remains relatively modest with chemoresistance (Tan et al., 2020). In highly proliferative tumors such as glioma, the increased metabolic rates lead to the accumulation of higher concentration of free radicals, leading to an increase in ROS levels (Olivier et al., 2020). However, the regulation of ROS is often known to maintain cellular stability and enhance tumor progression by modulating various proliferative pathways such as PI3K/AKT/mTOR pathway, MAPK pathway and Wnt signalling pathway (Precilla et al., 2021). Among these, the PI3K/AKT/mTOR pathway has been reported to be aberrantly activated in almost 80% glioma cases (Cancer Genome Atlas Research Network, 2008; Cheng et al., 2009). In addition, previous studies have demonstrated that low levels of ROS could promote glioma progression and are responsible for T resistance (Singh et al., 2021). Thereof, designing targeted therapies that can enhance the levels of ROS and impede glioma proliferation via inhibition of PI3K/AKT/mTOR pathway are vitally important as they might improve glioma patient therapeutic outcome (Taylor et al., 2019).

In this context, employing drug repurposing approach, we had attempted to trigger apoptosis in glioma via ROS-mediated inactivation of PI3K/AKT/mTOR pathway (Yang et al., 2020). This was on grounds of our previous study, which proved that M and E synergistically enhanced the anti-glioma efficacy of T on C6 murine glioma and U-87 MG human glioblastoma cells by alleviating oxidative stress (Kuduvalli et al., 2021). Underpinning this idea, the current study was carried out to gain insight if the triple-drug combination (TME) could induce oxidative stress-mediated deactivation of PI3K/AKT/mTOR pathway and promote apoptosis *in vivo.* In fact, individually T, M and E were reported to inhibit tumor cell growth, proliferation and promote apoptosis in various tumor types, including glioma (Sharifi-Rad et al., 2020).

It has been reported in previous literature that the combination of T and M significantly increased survival rates in animal models of glioma and in clinical setup (Seliger et al., 2019; Valtorta et al., 2021). Also, monotherapy with E in a preclinical orthotopic nude glioma-induced mice model had significantly increased the survival rates of the tumor-induced mice (Chen et al., 2011). In accordance with those findings, the survival benefits of our triple-drug combination treatment strategy on the established glioma-induced orthotopic xenograft model exhibited prolonged survival rates as compared to the tumor-control (**Figure 1B**).

Pathologically, glioma is characterized by hyper cellularity and haemorrhage, wherein a highly condensed cellular aggregation is a hallmark of glioma prognosis (Brat et al., 2002). Following the assessment of survival benefit, histopathological observation was carried out to analyse the effect of our drugs both individually and in combination on glioma tumor sections by H & E staining. Interestingly, in our study, glioma-induced rats treated with T-alone and M-alone was effective in inhibiting the tumor growth as individual regime; However, E-alone did not exhibit much effect as an individual drug. Nevertheless, combined administration of these drugs as a triple-drug combination was able to suppress the growth of glioma as evident by the reduction in haemorrhage that was almost similar to that of the normal control rats (**Figure 1C**). One attributed fact for this effect would be partially covered by the synergism of M and E along with clinically acceptable dose of T (10 mg/kg) which could have enhanced the inhibitory effect of T than their individual and dual counterparts.

Since, glioma is a highly proliferative and neovascularized tumor, combining two agents with two different effects along with a standard drug would help in synergistically regulating both proliferation and vascularization at a low dosage (Daisy Precilla et al., 2022). To be more specific, in highly vascularized tumors like glioma, there is high proliferation and extreme infiltration into surrounding tissues which is another target to be considered (S. K. Tan et al., 2018). Interestingly, in addition to reducing the population of proliferating glioma cells as evident from H & E, the triple-drug combination depicted a significantly reduced angiogenesis and proliferation as evident from the IHC and qRT-PCR analysis of Ki67, VEGF and its receptor VEGFR in tumor sections (**Figures 2 & 3B**). These results were in accordance with previous reports wherein the drugs, T and M were found to suppress glioma proliferation and neo-vascularization individually (Vasilev et al., 2018).

As glioma is a highly vascularised tumor, angiogenesis is vital for cell survival and tumor progression. One of the major contributing factors to angiogenesis and neovascularization is regulation and maintenance of ROS at low levels. However, increase in the levels of ROS leads to oxidative stress that is known to induce DNA damage (Ahir et al., 2020). Oxidative stress crops up when there occurs a mismatch between the antioxidant defence system in our body and an increase in the rate of reactive species production. This so-called oxidative therapy, induces the production of ROS, thereby triggering them towards apoptosis (Arfin et al., 2021). In this context, we aimed to determine the effect of the above-mentioned regimes on oxidative stress. The extent of the ROS production as a result of oxidative stress, is widely determined by DCF-DA. Generally, DCF-DA is reduced by esterase to DCF-H which is the further reduced to DCF by ROS (Wang and Roper, 2014). Elevated ROS levels are often known to activate the antioxidant and non-antioxidant defence system molecules, such as SOD, CAT, GPx, GSH and the marker for lipid peroxidation, TBARS. Among these, SOD and CAT are involved in the reclamation of superoxide molecules (O_2_^-^) and peroxides (H_2_O_2_) respectively. GPx, on the other hand, is involved in elimination of free radicals and peroxides and helps to maintain a stable redox homeostasis. (B. L. Tan et al., 2018) Similarly, TBARS is activated as a result of lipid peroxidation (Hasanuzzaman et al., 2020). In accordance with this idea, the triple-drug combination significantly enhanced the levels of ROS as evident by the DCF-DA assay (**Figure 3A**). In concurrence with our ROS activity, the levels of antioxidant and non-antioxidant enzymes were upregulated in the triple-drug (TME), dual-drug combination (TE) and individual treatment (E). One interesting fact to be noted is that, E, despite being an anti-oxidant, had elevated the levels of ROS, when administered at a higher concentration (150 mg/kg). This aspect unveils the ability of this drug to modulate its activity as an anti-tumor agent, which corroborated with the previous findings (Ouyang et al., 2020). However, profound investigation on this specific property of E is warranted in future.

With regard to regulation of oxidative stress in glioma, besides the aforementioned antioxidant and non-antioxidant enzymes, Nrf2 and HIF-1α are few among the validated targets for therapy (Godoy et al., 2020). Up-regulation of Nrf2 has known to enhance glioma proliferation by reversing the levels of ROS, leading to tumor cell survival (Wu et al., 2019). Hence, a deeper knowledge on Nrf2 pathway appeals a promising approach to overcome drug resistance in glioma. Also, as the octopus tumor proliferates, it outgrows existing vasculature and creates regions of hypoxia throughout the tumor. To overcome this hypoxia-induced stress, tumor cells employ the transcriptional regulator, HIF-1α (Kaur et al., 2005). HIF-1α promotes angiogenesis that induces glycolysis and switches off oxidative phosphorylation (Corcoran and O’Neill, n.d.; Nagao et al., 2019). This in turn rescues the tumor cells from hypoxic stress, and enhancing glioma stemness, invasiveness and chemoresistance (Wang et al., 2017). Growing evidences emphasize that Nrf2 and HIF-1α, both promotes chemo-resistance and contributes to carcinogenesis (Telkoparan-Akillilar et al., 2021). Therefore, developing inhibitors that targets Nrf2 and HIF-1α would be an effective therapeutic modality to combat glioma. Triple-drug combination therapy, in the present study revealed a decrease in the gene and protein levels of Nrf2 and HIF-1α (**Figure 3C**). One interesting observation from our study was that the triple-drug combination up-regulated the levels of ROS, which was indirectly proportional to the levels of Nrf2 and HIF-1α, that could impede the survival of glioma tumor. These results were in concordance with the previous findings (Fan et al., 2017).

Nrf2 and HIF-1α are known to regulate various metabolic and proliferative pathways, which are responsible for angiogenesis, tumor proliferation and chemoresistance. One such pathway is PI3K/AKT/mTOR pathway, a key regulator for glioma survival and proliferation, which has demonstrated to enhance oxidative stress, thereby promoting tumor cell survival in several tumors (Dong et al., 2021). Among the major drivers of this pathway, AKT serves as a central node for PI3K signalling, while PTEN acts as an antagonist of this pathway (Haddadi et al., 2018). Ligand binding and phosphorylation of RTKs such as VEGFR leads to activation of PI3K which in turn activates PDK1 and AKT1 through Phosphatidylinositol 4,5-bisphosphate-2 (PIP2) and PIP3 cascade (Sugiyama et al., 2019). Upon activation, AKT phosphorylates and activates a plethora of substrates involved in the regulation of apoptosis, cell cycle, glucose metabolism, tumor invasion and so forth (Li et al., 2016). Increased expression of PTEN antagonizes the P13K pathway by dephosphorylation of PIP3 to PIP2, preventing the activation of AKT. Reduced AKT in turn, reduces the expression of GSK-3β, and downregulates several target genes involved in cell proliferation leading to cell cycle arrest (Duda et al., 2020). Thereof, anti-cancer drugs targeting P13K/AKT/mTOR would inhibit glioma proliferation, angiogenesis and promote apoptosis. Indeed, this statement was proven in our study, wherein the combination of TME significantly reduced the levels of VEGFR, VEGF, PI3K, PDK1, pAKT1, GSK-3β, p-mTOR while the levels of PTEN was significantly enhanced, with an arrest at G1 phase of the cell cycle (**Figures 3C & D; Figure 4 A-D**). However, further research is required to elucidate the exact mechanism by which the triple-drug combination exerts this action and bring about cell cycle arrest.

Additionally, while speculating whether oxidative stress-induced inactivation of PI3K/AKT/mTOR pathway has led to apoptosis, the triple-drug combination triggered the percentage of apoptotic cells when compared with the tumor-control group (**Figures 4E & F**). The induction of this apoptotic effect was more deeply clarified by the assessment of major markers of apoptosis such as BCL2, BAX, BAD, caspase-9 caspase-8 and caspase-3 (**Figures 3E-H**) (Wang and Youle, 2009). BCL2, a class of anti-apoptotic proteins, modulates apoptosis and promotes survival in tumor cells (Carrington et al., 2017); While, the pro-apoptotic proteins BAX, and BAD, residing on the mitochondrial membrane is usually down-regulated in tumor cells (Garrido et al., 2006). Imbalance between these two protein families results in evasion of apoptosis in glioma (Wong, 2011). In this regard, the triple-drug combination attenuated BCL2 and elevated BAX and BAD expression. This may be due to the alterations in mitochondrial permeability that promotes cytochrome-C efflux (cyt-C); which in turn activates the downstream initiator caspase such as caspase-9 and caspase-8. The initiator caspases finally lead to the activation of the executioner caspase, caspase-3, thereby bringing about programmed cell death. Therefore, we speculate that the TME combinatorial treatment used in this study had potentially hampered glioma proliferation by ROS-induced inactivation of PI3K/ATK/mTOR pathway on induction of apoptosis.

Following assessment of the anti-tumor efficacy of our triple-drug combination in an *in vivo* orthotopic human glioma model, an attempt was also made to computationally predict the efficacy of these available drugs and their metabolites against human ortholog of VEGFR (3HNG). As discussed earlier, VEGF being the key mediator of angiogenesis and PI3K/AKT/mTOR pathway in glioma (Colardo et al., 2021; Karar and Maity, 2011), pre-clinical screening of a potent inhibitor that can block the active site of human VEGFR could be a preliminary step in the drug screening pipeline against glioma progression stimulated through angiogenesis. From our molecular docking studies, we found that T with a glide score of -7.938 kcal/mol demonstrated a potential inhibitory property against VEGFR, while other chosen drugs had also displayed a relatively modest inhibitory property, possibly due to the presence of imidazole ring. Imidazole has been demonstrated to possess broad range of chemical, and biological property (Rulhania et al., 2021) and proved to be an effective molecule in anti-cancer drugs (Ali et al., 2017). This ancillary analysis provides insights on the efficacy of triple drug combination by virtue of its protein-ligand interaction profile of various key drug molecules against VEGFR (**Figure 5 & table 2**). It may also be hypothesized that the effect of triple-drug combination could have been attributed by their difference in molecular interaction which is evident from the docking analysis. Taken together, the triple-drug combination may possibly evolve as a potential human VEGFR inhibitor cocktail for glioma patients in future.

**Table 2:** Interaction profiles of the experimental compounds with 3HNG

| Sl. No | Drugs | PubChem ID | Description | Glide Score kcal/mol | Interactions |
| --- | --- | --- | --- | --- | --- |
| 1 | T | 5394 | 3-methyl-4-oxoimidazo[5,1-d][1,2,3,5] tetrazine-8-carboxamide | -7.938 | 2 H- interactions with Csy192 and Asp1040 |
| 2 | MTIC | 54422836 | 3-methyl-(triazene-1-yl)imidazole-4-carboxamide | -6.829 | 2 H- interactions with Csy192 and Asp1040<br>π-π interaction with Phe1090 |
| 3 | AIC | 9679 | 4-amino-1H-imidazole-5-carboxamide | -5.736 | 2 H- interactions with Val907 and Glu878<br>π-π interaction with Lys816 |
| 4 | M | 14219 | 3-(diaminomethylidene)-1,1-dimethylguanidine; hydrochloride | -4.194 | 2 H- interactions with Glu878 and Asp1040 |
| 5 | GUA | 8859 | di-amino methylidene-eurea | -3.855 | 2 H- interactions with Glu878 and Ile1038 |
| 6 | E | 65064 | [(2R,3R)-5,7-dihydroxy-2-(3,4,5-trihydroxyphenyl)-3,4-dihydro-2H-chromen-3-yl] 3,4,5-trihydroxybenzoate | -3.397 | 3 H- interactions with Asp1040, Arg1021 and Asp807<br>π-π interaction with His1020 |

**Table 3:** Highest and lowest occupied molecular orbitals’ energies

| Sl. No. | Drug | HOMO | LUMO | Energy gap |
| --- | --- | --- | --- | --- |
| 1 | T | -0.2522 | -0.0903 | -0.16194 |
| 2 | MTIC | -0.2112 | -0.0525 | -0.15871 |
| 3 | AIC | -0.1873 | -0.0072 | -0.18009 |
| 4 | M | -0.3975 | -0.1617 | -0.23585 |
| 5 | GUA | -0.4416 | -0.1825 | -0.25915 |
| 6 | E | -0.2025 | -0.0303 | -0.17217 |

In all the cases, the HOMO-LUMO gap is minimal with 0.19 eV being the average energy difference indicating the fragility of the bound electrons and molecular reactivity (Banavath et al., 2014; Selvaraj et al., 2021b). 54422836 has the lowest HOMO LUMO energy gap of 0.15 eV while 8859 had the highest HOMO-LUMO gap of 0.26 eV. The higher HOMO energy compared to the LUMO energy of all these reported molecules indicates a better electron donating ability than electron accepting ability. The high electron donating ability could have attributed to the overall better affinity of these reported molecules to the target protein.

However, there are a few limitations in the current study. A xenograft rat model was employed in this study; instead, immune-compromised models are warranted for further analysis of this particular drug combination in glioma.

In conclusion, while the search for new compounds to manage glioma are in row, there is a pressing need to mitigate the increasing prevalence of glioma. Data from our study showed that the combination of the existing drugs, M and E synergistically in enhanced the anti -glioma potency of T in a preclinical orthotopic xenograft glioma model. Further, it was also found that the mechanism of inhibiting glioma proliferation was attributed to the induction of apoptosis via ROS-mediated inactivation of PI3K signaling pathway. Hence, the combination of TME could serve as a potential therapeutic agent for the treatment of glioma patients in the near future. Nevertheless, future clinical validations are required to evaluate the therapeutic efficacy of the combination of TME among glioma patients.

## 5. Author contribution statement

**Anitha T.S:** Conceptualization, Resources, Funding Acquisition, Methodology, Supervision, Writing-Review & Editing. **Shreyas S Kuduvalli:** Data Curation, Writing-Original draft preparation, Writing-Review & Editing. **Daisy Precilla S**: Data Curation, Writing-Review & Editing**. Anandraj Vaithy:** Data Curation. **Mugilarasi Purushothaman:** Investigation, Formal analysis & Data Curation. **Muralidharan Arumugam Ramachandran:** Data Curation & Software. **Agiesh Kumar B:** Visualization & Investigation. **Markus Mezger:** Visualization. **Justin S Antony**: Formal analysis. **Madhu Subramani:** Supervision. **Biswajit Dubashi:** Project Administration. **Indrani Biswas:** Data Curation. **K P Guruprasad:** Supervision.

## 6. Acknowledgments

We would like to thank Prof. Adithan C, Former Dean-Research, Sri Balaji Vidyapeeth for providing Dr. Vany Adithan Research Fellowship for the first author. We would also like to acknowledge Prof. K Satyamoorthy, Director, Manipal School of Life Sciences, MAHE, Manipal, for providing the animal facility to carry out our objective. We acknowledge Centre for Stem Cell Research (a unit of inStem, Bengaluru), CMC Campus, Vellore, India for providing flow-cytometry facility. We express our gratitude towards Sri Balaji Vidyapeeth for providing the basic laboratory and instrumentation facilities. We would also like to thank Mr. Guillermo Urena-Bailen for the graphical illustration.

## 7. Contribution to the Field Statement

While the search for new compounds to manage glioma are in row, there is a pressing need to mitigate the increasing prevalence of glioma. Repurposing of drugs has emerged as promising strategy, especially when the usual anti-cancer monotherapy has demonstrated failures in the safety and tolerability of oncological patients. Accordingly, data from this study showed that this novel cocktail of repurposed drugs, metformin and epigallocatechin gallate displayed synergism in enhancing the anti-glioma potency of temozolomide in a preclinical orthotopic xenograft glioma model. Further, it was also found that the mechanism of inhibiting glioma proliferation was attributed to the induction of apoptosis via ROS-mediated inactivation of PI3K signalling pathway. Hence, this novel cocktail of temozolomide, metformin and epigallocatechin gallate could serve as a potential therapeutic agent for the treatment of glioma patients in the near future. Nevertheless, future clinical validations are required to evaluate the therapeutic efficacy of this combination among glioma patients.

## 10. Data Availability Statement

The datasets generated or analysed during the current study are available from the corresponding author on reasonable request.

## Abbreviations

C: Control
TC: Tumor-control
T: Temozolomide
M: Metformin
E: Epigallocatechin gallate
TM: Temozolomide + Metformin
TE: Temozolomide + Epigallocatechin gallate
ME: Metformin + Epigallocatechin gallate
TME: Temozolomide + Metformin + Epigallocatechin gallate
MTIC: 3-methyl-(triazen-1-yl) imidazole-4-carboxamide
AIC: 4-amino-1H-imidazole-5-carboxamide
GUA: Guanylurea

