## Supplementary Table 1 for "A combination of Metformin and Epigallocatechin Gallate Potentiates Glioma Chemotherapy *in vivo*"

**Supplementary Table S1** Primer sequences used for the qRT-PCR analysis, that were designed using Primer BLAST, NCBI.

| Gene | Forward (5'-3') | Reverse (5'-3') | ID |
|---|---|---|---|
| **β-Actin** | AGATGACCCAGATCATG TTTGAGA | GCATGAGGGAGCGCGT AA | NM_031144.3 |
| **Nrf2** | CAGCATGATGGACTTGG AATTG | GCAAGCGACTCATGGT CATC | NM_031789.2 |
| **HIF-1α** | GCAACTAGGAACCCGAA CCA | TCGACGTTCGGAACTC ATCC | NM_024359.2 |
| **VEGF** | CAAACCTCACCAAAGCC AGC | ACGCGAGTCTGTGTTTT TGC | NM_031836.3 |
| **VEGFR-1 (FLT-1)** | CAGTGGCTCCACGACCT TAG | GGTGAGGTACGCTGAG CTTT | NM_019306.2 |
| **PI3K** | TGGCCCGGGTAGGTTTG AAT | ATGCCCTAGGTGACCT GACA | NM_00137130 0.2 |
| **PTEN** | AAAGCTGGGAAAGGACG GAC | CACCTTTAGCTGGCAG ACCA | NM_031606.1 |
| **PDK1** | GGCATAGAGCGGCAGGT TG | AGAAGCGCGCATAGAA GTCC | NM_053826.2 |
| **AKT1** | TCATTGAGCGCACCTTCC AT | TTCTGCAGGACACGGTT CTC | NM_033230.3 |
| **mTOR** | CTGCACTTGTTGTTGCCT CC | ATCTCCCTGGCTGCTCC TTA | NM_019906.2 |
| **GSK3-β** | AACTCCACCAGAGGCAA TCG | AAGCGGCGTTATTGGT CTGT | NM_032080.1 |
| **Bax** | AGACACCTGAGCTGACC TTGG | GTTGTTGTCCAGTTCAT CGCC | NM_017059.2 |
| **Bcl2** | GGTGAACTGGGGGAGGA TTG | AGAGCGATGTTGTCCA CCAG | NM_016993.2 |
| **BAD** | CTAGGCTTGAGGAAGTC CGAT | CGGGAATGTGGAGCAG ATCA | AF279911.1 |
| **Caspase-8** | TTTCCATATCAGTCGGCG GG | TCAAGCAGGCTCGAGT TGTC | NM_022277.1 |
| **Caspase-9** | AGTTCCCGGGTGCTGTCT AT | GCCATGGTCTTTCTGCT CAC | AF271996.1 |
| **Caspase-3** | ATCCACGGAGGTTTCGT TGTTG | TGGGGCCAATAGTGTTT GGTA | NM_012922.2 |
