## Supplementary Figure 1 for "A combination of Metformin and Epigallocatechin Gallate Potentiates Glioma Chemotherapy *in vivo*"

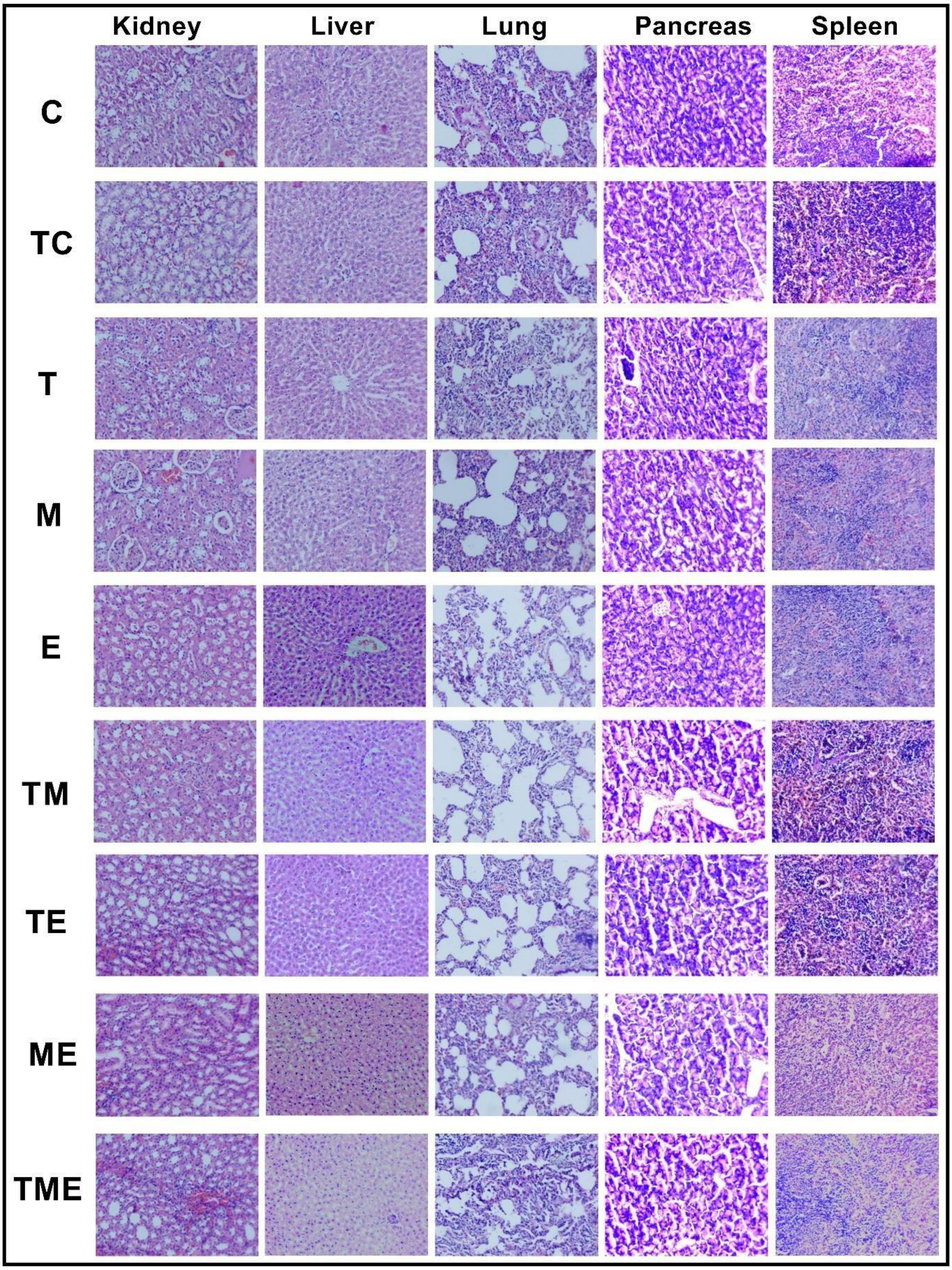


**Supplementary Figure S1:** Histopathological analysis of major organs (kidney, liver, lung, pancreas and spleen) of all the experimental groups with H&E stain.
