## Supplementary Figure 2 for "A combination of Metformin and Epigallocatechin Gallate Potentiates Glioma Chemotherapy *in vivo*"

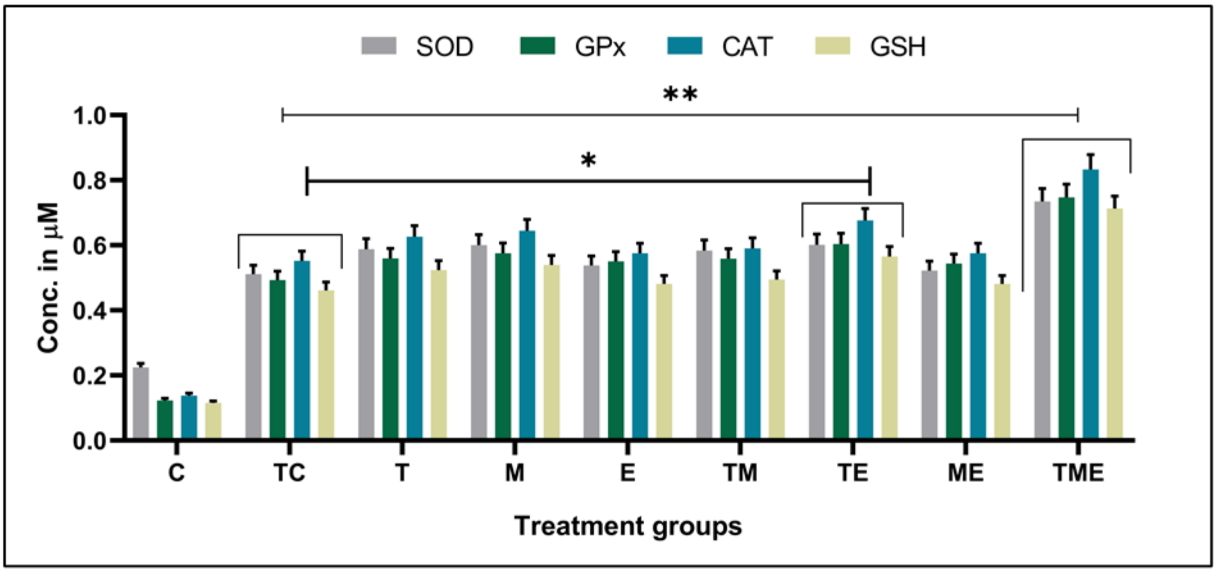


**Supplementary Figure S2:** Shows the levels of antioxidant and non-antioxidant enzymes (SOD, GPx, CAT, GSH) wherein, the triple-drug combination significantly enhanced the levels of all the enzymes followed by the dual-drug treatment (TE) (*P<0.01, ** P<0.001).
