## Supplementary Figure 3 for "A combination of Metformin and Epigallocatechin Gallate Potentiates Glioma Chemotherapy *in vivo*"

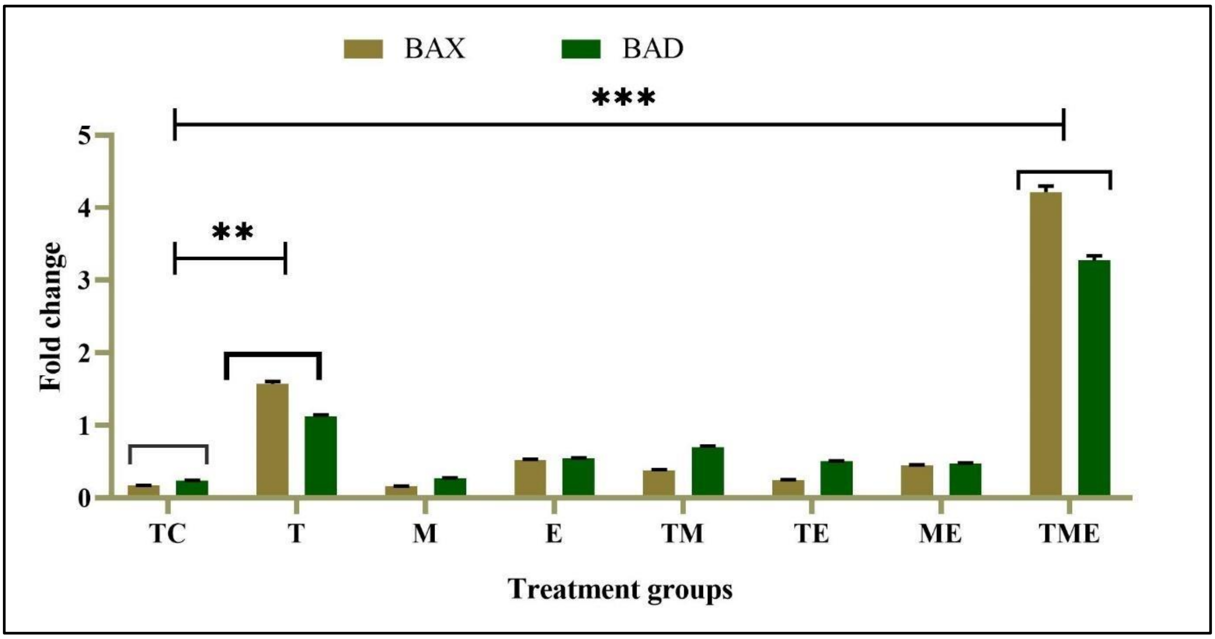


**Supplementary Figure S3:** Shows the gene expression levels of pro-apoptotic markers BAX and BAD, wherein, TME significantly elevated the levels of BAX and BAD, followed by the individual treatment with T (**P<0.01, *** P<0.001).
